# The large GTPase, mGBP-2, regulates Rho family GTPases to inhibit migration and invadosome formation in Triple-Negative Breast Cancer cells

**DOI:** 10.1101/679514

**Authors:** Geoffrey O. Nyabuto, John P. Wilson, Samantha A. Heilman, Ryan C. Kalb, Ankita V. Abnave, Deborah J. Vestal

## Abstract

Breast cancer is the most common cancer in women. Despite advances in early detection and treatment, it is predicted that over 40,000 women will die of breast cancer in 2019. This number of women is still too high. To lower this number, more information about the molecular players in breast cancer are needed. Several members of the Guanylate-Binding Proteins (GBPs) have been correlated with better prognosis in breast cancer. In this study, we asked if the expression of GBP-2 in breast cancer merely provided a biomarker for improved prognosis or whether it actually contributed to improving outcome. Specifically, we asked whether murine GBP-2 exhibited cell autonomous behavior that altered breast cancer cells in a manner predicted to improve prognosis. To answer this, the 4T1 model of murine Triple-Negative Breast Cancer was used. 4T1 cells themselves are highly aggressive and highly metastatic, while 67NR cells, isolated from the same tumor, do not leave the primary site. mGBP-2 did not alter cell proliferation in these two cell lines, but mGBP-2 expression inversely correlated with migration. More specifically, mGBP-2 inhibits cell migration and invadopodia formation by inhibiting the activation of Rac1, while promoting the activation of CDC42 and RhoA. Together these data demonstrate that mGBP-2 is responsible for cell autonomous activities that make breast cancer cells less aggressive.

## Introduction

Breast cancer strikes 1 in 8 women in the U.S, or about 12.3% of women (1). In 2015, over 242,000 cases of breast cancer were diagnosed in women and over 41,000 women died of the disease (2). Breast cancer remains the most common cancer in women and the second most deadly (2). While significant improvements have been achieved in early detection and the treatment of some types of breast cancer, as the numbers of older Americans increase so does the number of women with breast cancer (2). In addition, while the biomarkers and clinical targets in breast cancer have expanded, there are still some types of breast cancer, such as triple-negative breast cancer (TNBC), that lack targeted therapies and are short on biomarkers. Clearly we need more biomarkers to predict clinical outcomes and additional targets to improve therapy.

One family of proteins that has been implicated in predicting better clinical outcomes in breast cancer is the Guanylate-Binding Proteins (GBPs). The GBPs are a family of interferon- and cytokine-induced large GTPases (reviewed in (3)). Best studied for their anti-microbial activity, the GBPs are also implicated in cancer progression and prognosis. Human GBP-1 (hGBP-1) expression is correlated with better prognosis in breast and colorectal cancers (4-7). In breast cancer, hGBP-1 is part of a 5-gene signature that predicts with 85% accuracy a better than 10-year recurrence free survival (6). GBP-1 is expressed in both tumor and non-tumor cells in these tumors (6). This gene signature also includes the transcription factor, STAT1, which is responsible for the IFN-induction of GBPs. STAT-1 deficient mice develop breast cancer spontaneously (8). Recent gene array studies have identified two GBPs as highly expressed in a subtype of basal-like TNBCs with a greater than 80% recurrence free-survival for greater than 10 years but not in basal-like TNBCs with about a 35% 5-year survival (9). These were GBP-5 and GBP-1 (9). In that study the GBPs were expressed in a gene profile suggestive of an IFN-*γ* response and recruitment of B-, T-, and NK cells (9). Closely related human GBP-2 also correlates with improved metastasis-free interval in node negative breast cancers (10).

GBPs are some of the most abundant IFN-*γ*-induced proteins (11). Because the presence of GBPs in these breast cancers is associated with IFN-*γ*-induced response, it raises the question of whether GBPs are merely markers of an IFN-*γ* environment or whether these proteins play an active role in the improved prognosis. It is also unclear that even if GBP expression is important and consequential to the improved outcome, whether it is plays a significant role in the tumor cells, nontumor cells, or both. GBP-1 has been demonstrated to modulate T-cell antigen receptor signaling through its interactions with the cytoskeleton (12). It also inhibits the proliferation and migration/invasion of endothelial cells, thereby inhibiting neo-angiogenesis (13,14). GBP-1 also has properties that would suggest it could improve prognosis in a tumor cell-autonomous manner. GBP-1 inhibits epithelial cell and colorectal cancer cell proliferation (4,15). It also interacts with the actin cytoskeleton of HeLa cells and inhibits actin polymerization, thereby inhibiting migration (16). GBP-2 inhibits breast cancer cell invasion, in part, by inhibiting mitochondrial fission (17). Together these data suggest that GBPs could play roles in both tumor and non-tumor cells in breast cancer.

In this study, we examined the role of murine GBP-2 (mGBP-2) in the 4T1 model of murine breast cancer. We chose to look at the cell autonomous activity of mGBP-2 in these cells, with any eye toward later studies that would examine it’s extra-tumoral role. Like GBP-1 and GBP-2, mGBP-2 has been shown to influence cell proliferation and to inhibit cell migration (18,19). We show that mGBP-2 correlates with better prognosis in this model system and contributes to better outcome not by inhibiting cell proliferation but by significantly inhibiting cell migration and invadosome formation, as a consequence of or coincident with activating CDC42 and RhoA and inhibiting the activation of Rac1.

## Materials and Methods

### Cells and cell culture

4T1 and 67NR cells were the gift of Dr. Fred Miller (Karmanos Cancer Center, Wayne State University, Detroit, MI) (20). Cells were cultured in Dulbecco’s Modified Eagle’s medium (DMEM) with 4.5 g/L glucose (Mediatech, Manassas, VA), 10% fetal bovine serum (FBS; Atlanta Biologicals, Lawrenceville, GA), 50 µg/ml penicillin/streptomycin (Mediatech) and 2 mM L-glutamine (Mediatech) at 37°C in 5% CO_2_. Cells were suspended by treatment with 0.05% trypsin/0.53 mM EDTA (Mediatech).

### Western blots

Cells were lysed in RIPA (50 mM Tris, pH 7.5, 150 mM NaCl, 1% NP-40, 0.1% SDS, 0.5% sodium deoxycholate) containing 1 µl/ml Protease Inhibitor Cocktail (Sigma, St. Louis, MO), and 1 mM phenylmethanesulphonylfluoride (PMSF). Where appropriate phosphatase inhibitors have been added to the lysis buffer to a concentration of 25 mM sodium fluoride and 10 mM sodium vanadate. Protein concentrations were determined using BCA Protein Assay (Bio-Rad, Hercules, CA). Cell lysates were size fractionated on 8 or 15% SDS-PAGE gels using Tris-Glycine-SDS Tank buffer (25 mM Tris, 0.192 M Glycine, 0.1% SDS) and transferred to Immobilon-PVDF membrane (Millipore, Billerica MA). Wet transfer was performed using 1X transfer buffer (25 mM Tris, 0.192 M Glycine, 20% methanol) at 100 volts for 120 min. The membrane was blocked in TBST (250 mM Tris, pH 8, 5 M NaCl, 0.3% Tween-20) plus 5% nonfat dry milk for 1 hr at room temperature (RT) or overnight (ON) at 4°C. Membranes were incubated for 1 hr at RT or ON at 4°C in the primary antibodies described below. Membranes were washed in TBST three times for 15 min each. Membranes were washed three times for 15 min each in TBST. Chemiluminescence was detected using Super Signal West Pico Chemiluminescent Substrate (ThermoFisher Scientific, Rockford, IL). Where necessary, the membranes were stripped using Restore Western Blot Stripping Reagent (ThermoFisher Scientific).

### Antibodies

The following primary antibodies were diluted in TBST plus 5% nonfat dry milk for western blot: rabbit anti-mGBP-2 (1851 (21), 1:20000), rabbit anti-actin (A2066; 1:3000) (Sigma), rabbit anti-*α*-tubulin (GTX102078; 1:10000) (GeneTex, Irvine, CA), mouse anti-GAPDH (60004; 1:20000) (Proteintech, Rosemont, IL). The above antibodies were incubated with membrane for 1 hr at RT. The rabbit anti-RhoA (2117T; 1:4000) (Cell Signaling, Danvers, MA), mouse anti-Rac1 (610650; 1:4000) (BD Biosciences, San Jose, CA), mouse anti-CDC42 (610928; 1:250) (BD Biosciences), rabbit anti-Akt (9272; 1:3000) (Cell Signaling), and rabbit monoclonal anti-phosphoAkt (4060 (ser 473); 1:4000) (Cell Signaling) were incubated with membrane overnight at 4°C. Hrp-conjugated goat anti-mouse immunoglobulin (1:8000) or goat anti-rabbit immunoglobulin (1:6000) (Jackson ImmunoResearch, West Grove, PA) were diluted in TBST containing 5% w/v nonfat dry milk and incubated with the membranes for 1 hr at RT.

### Colony assay

Cells (1 × 10^3^ cells/6-cm dish) were seeded in triplicates. Whenever necessary, cells were treated with recombinant murine IFN-*γ* (IFN-*γ*; PBL Assay Science, Piscataway, NJ) at the concentrations listed. Media was changed every two days and after culture, cells were washed in 2 ml PBS (138 mM NaCl, 2.6 mM KCl, 5.4 mM Na_2_HPO_4_, 1.8 mM KH_2_PO_4_, pH 7.4), fixed in ice-cold 100% methanol for 5 min, and stained with 1% w/v crystal violet (ThermoFisher Scientifics) for 5 min. All the colonies per dish with ≥ 50 cells per condition were counted.

### Click-It EdU

Cells were plated on 22-mm coverslips in 6-well dishes. Whenever necessary, cells were treated with rmIFN-*γ* (0, 100, 250, and 500 U/ml) in duplicates. Cells were incubated with 20 uM 5-ethynyl-2’-deoxyuridine (EdU) (Carbosynth, San Diego, CA) for 30 min, fixed with 4% paraformaldehyde for 10 min at RT, permeabilized with 0.2% Triton X-100 for 10 min at RT, washed with PBS and incubated with 200 μl labelling mix (2mM CuSO_4_.5H_2_O, 10 μM Cyanine5 azide (Lumiprobe, Hunt Valley, MD) and 20 mg/ml Ascorbic acid) for 30 min at RT. Cells were washed with PBS and stained with 150 nM DAPI for 5 min at RT. Coverslips were mounted with Fluoromount-G (SouthernBotech, Birmingham, AL) (22). Four random fields per coverslip were imaged at 20X on a Cytation 5 Imaging Multi-Mode Reader (BioTek Instrument, Winooski, VT) using DAPI and Texas red filters. Cells from the Cytation 5 images were manually counted using ImageJ software (National Institutes of Health, Bethesda, MD, USA) and the percentage of EdU positive cells (red) per field was calculated.

### Boyden chamber

Whenever necessary, cells were pre-treated with 100 U/ml IFN-*γ* for 24 hrs. Cells (1 × 10^4^) were seeded on 8 µm Boyden chamber inserts (Becton Dickinson Labware, Bedford, MA)(23) coated with fibronectin (R&D Systems, Minneapolis, MN) (5 μg/ml) on both sides. Three hundred microliter of serum-free media (SFM) was added to the inserts and the bottom wells of 24-well plates with or without 100 U/ml IFN-*γ*. Cells were allowed to adhere for 3 hrs. The SFM in the bottom well was aspirated and 500 μl 20% FBS in DMEM was added to the bottom chamber. After 5 hrs, unmigrated cells were removed from the inside of the inserts with a cotton swab and the cells were fixed in ice-cold 100% methanol for 5 min and stained with 1% w/v crystal violet (ThermoFisher Scientifics) for 3 min. The entire membrane was then imaged at 4X on a Cytation 5 Imaging Multi-Mode Reader. All migrated cells were counted manually using ImageJ cell counter software.

### Wound healing assay

Cells were plated in duplicates in 6-well dishes and incubated for 24 hrs at 37°C to confluence. A 200 μl pipette tip was used to create a scratch and the debris was removed with a PBS wash. Three fields per well were photographed immediately (0 hr) and 24 hr after scratch formation on an EVOS FL Inverted Microscope (Thermofisher) at 4X (24). The wound width at 0 hr and 24 hrs post-introduction of scratch was measured using MetaMorph (Molecular Devices, San Jose, CA). The difference between wound width at 24 hrs and at 0 hr was used to calculate the percent of the wound closure in relation to wound width at 0 hr.

### Generation of control and mGBP-2-directed shRNAs

pSIH-H1 was used for the generation of shRNAs against eGFP and mGBP-2, following the instructions from System Biosciences (Mountain View, CA). Briefly, both top strand and complementary strand oligonucleotides were generated that when allowed to anneal contained BamH1 and EcoR1 overhangs that were complementary to the overhangs in the pSIH-H1 vector. This provided directional cloning of the resulting shRNA into the vector. The top and complementary strand oligonucleotides for each shRNA were first allowed to anneal. To phosphorylate the insert, a 20 µl mixture was set up with 1 µM each of the top and complementary oligonucleotides, 1 mM ATP, 1X T4 kinase buffer, and 20 units of T4 polynucleotide kinase. Using a thermocycler, the reaction was heated to 37°C for 30 min, followed by 95°C. After 2 minutes, the machine was turned off and the samples were allowed to cool to RT. The resulting double stranded oligonucleotides were then ligated into BamH1/EcoR1 cut and purified pSIH-H1. Ligated plasmid was used to transform DH5*α E.coli* cells per manufacturer’s instructions (Invitrogen). Colonies were isolated from LB plates containing 50 µg/ml ampicillin and at least 10 colonies per construct were grown in 100 µl of LB containing ampicillin at 37 °C ON. Those plasmids containing inserts were identified by PCR amplification from the liquid cultures following manufacturer’s instructions. Samples were then separated on 3% agarose gels in 1X TAE and the presence of an insert of 105 bp indicated the presence of an insert. The sequencing primers were: 5’-TTAGCCAGAGAGCTCCCAGGCTCAGA-3’ for forward and 5’-TCACCATAAACGTGAAATGTCTTT-3’ for reverse. Two shRNAs were generated against eGFP. FOR eGFP shRNA #2 the top strand was 5’-GATCCCACAAGCTGGAGTACAACTACAACAGCCACTTCCTGTCAGATGGCTGTTGTAGTTGTACTCCAGCTTGTGTTTTTG-3’ and the complementary strand was 5’-AATTCAAAAACACAAGCTGGAGTACAACTACAACAGCCATCTGACAGGAAGTGGCTGTTGTAGTTGTACTCCAGCTTGTGG-3’. For mGBP-2 shRNA #3 the forward strand was 5’-GATCGATGTTGTTGAAACACTTCTACTCGAGTAGAAGTGTTTCAACAACATCTTTTTG -3’ and the complementary strand was 5’-AATTCAAAAAGATGTTGTTGAAACACTTCTACTCGAGTAGAAGTGTTTCAACAACATC-3’.

### Transfection and cell selection

4T1 and 67NR cells at 70-80% confluence were transfected with the shRNA constructs described above in FuGene 6 at a ratio of 3:2 per manufacturer’s instructions (Thermo Fisher). Pools of transfected cells were selected in media with 7 µg/ml puromycin. After selection, the cells were maintained in media containing 5 µg/ml puromycin.

### Immunofluorescence

Cells (4 × 10^4^) were plated on 12-mm coverslips in duplicates in 24-well dishes and serum starved for 3 hrs before adding warm 10% FBS in DMEM for 20 min. Cells were fixed with 4% paraformaldehyde for 10 min at RT, permeabilized with 0.2% Triton X-100 for 10 min at RT, and washed with PBS. The cells were blocked with 0.5 ml antibody (Ab) dilution buffer (PBS containing 0.05% Tween 20, 3% BSA, 5% glycine) with 10% nonimmune horse serum at room temperature for 1hr. Next, 100 μl of diluted Alexa fluor 594 phalloidin (Molecular Probes, Eugene, OR) in Ab dilution buffer with no horse serum (1:200) was added and incubated overnight. Cells were washed with PBS and stained with 150 nM DAPI for 5 min at RT. Coverslips were mounted with Permount-G (SouthernBotech). Random fields per coverslip were imaged at 20X and 40X on an EVOS FL Inverted Microscope (Thermofisher) using DAPI and Texas Red filters. The images were loaded onto ImageJ software and 100 cells per condition were used to measure parameters that included surface areas, elongation ratios, number of projections and length of projections.

### Image Analysis

To measure cell surface areas and elongation indices, images were uploaded onto ImageJ and converted to 8 bit. Images were subjected to percentile threshold algorithm. After applying the threshold settings, the “analyze particle” function was used with the pixel size (pixel^2^) set from 0–∞ and circularity set from 0–1.0 to include all particles. Results from individualized cells were determined. The data output included object count (# of individual objects), total area of detected objects (total pixels^2^), and aspect ratio (elongation index) (25). The cell surface area was converted to μm^2^ by dividing the total pixels^2^ by 9.61 (1 μm = 3.1 pixels). To determine the number of cell projections and length, the images were uploaded onto ImageJ. The projections were counted visually and the lengths were measured by drawing a straight line along the cell projections.

### Rho GTPase activity assays

Cells were plated in 15-cm dishes to confluence and serum starved overnight. Cells were then incubated with 15 ml 20% FBS in DMEM for 30 mins, washed with 5 ml ice-cold PBS + 1 mM MgCl_2_ and lysed in 1ml of either ice-cold Pak-Binding Domain (PBD) buffer (for Rac1 and Cdc42 pull downs) (50 mM Tris pH 7.4, 150 mM NaCl, 10 mM MgCl_2_, 1% Triton X-100, 1 mM phenylmethylsulfonyl fluoride (PMSF), 10 µl/ml protease inhibitors.) or ice-cold Rhotekin-Binding Domain (RBD) buffer (for RhoA pull downs) (50 mM Tris-HCl (pH 7.4), 500 mM NaCl, 1% (vol/vol) Triton X-100, 0.1% (wt/vol) SDS, 0.5% (wt/vol) deoxycholate and 10 mM MgCl_2,_ 1 mM PMSF, 10 µl/ml protease inhibitors). Cells were scraped, collected in a microcentrifuge, and vortexed briefly. The lysates were centrifuged at 14,000 RPM for 3 min at 4 °C in a tabletop centrifuge. The supernatant was transferred to a fresh tube and snap frozen in liquid nitrogen. The protein concentrations were determined using the Bio-Rad DC protein assay kit. One mg-1.5 mg of protein was brought to 1 ml volume using either PBD or RBD lysis buffer. GST-Rhotekin-RBD or GST-PAK-PBD beads (50 µg) was added to the lysate samples and rotated for 45 min at 4 °C, centrifuged at 12, 000 RPM for 3 min and the supernatant discarded using a 27.5-gauge needle. The beads were washed three times with 1 ml PBD wash buffer (50 mM Tris pH 7.6, 150 mM NaCl, 1 % Triton X-100, 10 mM MgCl_2_) and resuspended in 30 µl total cell lysates were run in a 15% SDS-PAGE gels, transferred, and probed as previously described. The scanned X-ray films were uploaded onto ImageJ to measure the densitometric values of active and total GTPases. The ratio of active and total GTPases densitometric values were calculated and represented on the graph as the fold of active GTPase ± S.D relative to control 67NR cell line, which was assigned an arbitrary value of 1.

### Gene expression profiling and data processing

The program Km plot (26) was used to analyze the data from the following publically available microarray data sets: E-MTAB-365 (n=537), E-TABM-43 (n=37), GSE11121 (n=200), GSE12093 (n=136), GSE12276 (n=204), GSE1456 (n=159), GSE16391 (n=55), GSE16446 (n=120), GSE16716 (n=47), GSE177705 (n=196), GSE17907 (n=54), GSE18728 (n=61), GSE19615 (n=115), GSE20194 (n=45), GSE20271 (n=96), GSE2034 (n=286), GSE20685 (n=327), GSE20711 (n=90), GSE21653 (n=240), GSE2603 (n=99), GSE26971 (n=276), GSE2990 (n=102), GSE31448 (n=71), GSE31519 (n=67), GSE32646 (n=115), GSE3494 (n=251), GSE37946 (n=41), GSE41998 (n=279), GSE42568 (n=121), GSE45255 (n=139), GSE4611 (n=153), GSE5327 (n=58), GSE6532 (n=82), GSE7390 (n=198), and GSE9195 (n=77). The Affymetrix probe ID for GBP-2 used for the analysis was 242907_at. For gene array analyses of all breast cancers, there was no filtering for hormone status, intrinsic subtype (histology), grade, lymph node status, or treatment. For gene array analysis of TNBC tumors, only the ER negative, PR negative, and unamplified HER2 tumors were analyzed. Again, no filter for intrinsic subtype, grade, or treatment was employed.

The patients were split by the median value into low versus high expression. The RNAseq ID was gbp2. Analysis was not restricted to stage, gender, grade, or race.

### Invadopodia analysis

Cells on coverslips were treated with 1 µM Phorbol 12,13-dibutyrate (PDBu; Sigma) for 30 minutes. After fixation with 3.7% paraformaldehyde and permeabilization with 0.1% Triton X-100, the cells were stained with Alexa Fluor 488 phalloidin (1:50; Molecular Probes) and anti-cortactin (1:450; p80/90 clone 4F11; 05-180; Millipore) for 1 hour at room temperature. After incubation with Alexa Fluor 594 anti-mouse for 45 minutes, the cells were stained with 150 nM DAPI before mounting in Permount G. Random fields per coverslip were imaged at 60X oil on an EVOS FL Inverted Microscope using GFP and Texas Red filters. At least 50 cells from each cell line were examined for invadosomes in each experiment and the data is presented as average percentage of cells containing invadopodia ± standard deviation.

### Statistical analysis

Statistical analyses were carried out by two-tailed t-tests when two groups were analyzed. One-way Anova (GraphPad Software, La Jolla, CA) was utilized when more than two groups were analyzed and subsequent comparisons were done by Tukey’s post-test. Statistically different groups are defined as * P < 0.05, ** P < 0.01, *** P < 0.001 and **** P < 0.0001.

## Results

STAT1 is a transcription factor responsible for GBP expression downstream of interferon exposure. Mice lacking STAT1 spontaneously develop mammary carcinomas (8). Where the expression of hGBP-1 is part of a gene signature that correlates with improved prognosis of human breast cancers, hGBP-1 is expressed in both tumor cells and the surrounding stroma (6). Forced expression of hGBP-1 in a murine breast cancer cell line inhibited it proliferation (27). hGBP-2 is a single marker of improved metastasis-free intervals in node negative breast cancers and is proposed to correlate with tumor T cell responses (10). Finally, an extensive analysis of microarray data in breast cancers indicates that hGBP-1 and hGBP-5 are robustly induced in a subset of basal-like triple negative breast cancers with greater than 80% recurrence free survival for 10 years (9). These tumors also had a robust gene signature of active B-, T-, and NK cells, suggesting a robust immune cell infiltration (9).

To expand on our understanding of the role of GBP-2 in human breast cancer, we asked whether GBP-2 correlated with improved recurrence-free survival (RFS) and/or overall survival (OS) in a series of gene array studies of all breast cancer types, hormone status, grades, and node status (Figure 1). Combined data from over 1700 patients (883 with low and 881 with high GBP-2) showed that tumors expressing high levels of GBP-2 had significantly better RFS (Hazard ratio = 0.67 and 95% CI of 0.57 – 0.78) (Figure 1A). Analysis of correlation of GBP-2 expression and OS in 626 patients also showed that high GBP-2 expression correlated with improved OS for a group including all breast cancers (Hazard ratio = 0.63 and 95% CI of 0.46 – 0.86) (Figure 1B). The improved correlation with elevated GBP-2 expression from microarray data was confirmed with RNA seq. data for both RFS (Figure 1C) and OS (Figure 1D). Because the model of breast cancer that we will be using involves triple-negative breast cancers, we asked how GBP-2 expression correlates with RFS in TNBCs. Human GBP-2 correlates with improved RFS in human TNBCs of all grades, histological type, or node status (Figure 1E).

**Figure 1.**
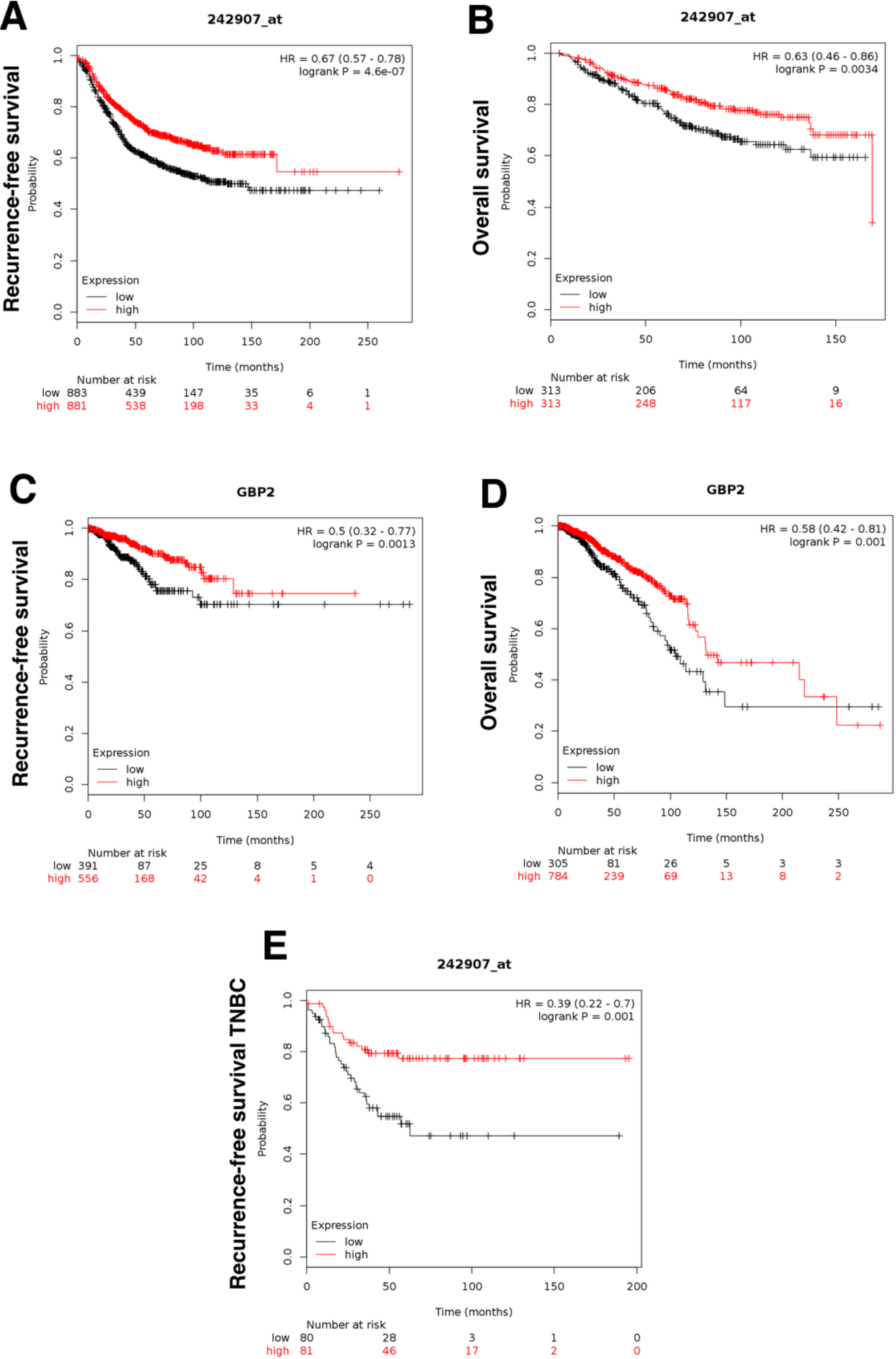
GBP-2 correlates with better recurrence-free (RFS) and overall survival (OS) in human breast cancers. **A).** The probability of RFS versus time for breast cancers of all types, stages, and grades was plotted for those tumors with high and low levels of GBP-2 expression. **B).** The OS of patients of patients with all subtypes, stages, and grades was plotted for those tumors with high versus low GBP-2 expression versus time. **C).** RNA seq data was used to confirm the array data for GBP-2 and RFS. **D).** RNA seq data was used to confirm the array data for GBP-2 and OS. **E).** Because the cell lines that we are using for this study are TNBCs, the correlation between GBP-2 expression and RFS in TNBC tumors was plotted.

These studies raise the question of what role(s) GBPs play to possibly be protective in breast cancer and whether these reflected the expression of GBPs in the tumor cells, stromal cells, or both. To examine the cell autonomous role of GBPs in breast cancer, the murine 4T1 model of metastatic breast cancer was chosen. This series of tumor cell lines were originally isolated from a mammary TNBC that arose spontaneously in a Balb/c mouse (20). Investigators generated sublines from the heterogenous murine breast cancer that differed in their ability to metastasize. This study focuses on two of these sublines: 4T1 and 67NR. 4T1 cells are highly aggressive, highly metastatic cells that rapidly metastasize to lungs and other organs. In contrast, 67NR cells are not metastatic and do not leave the primary site after injection into murine mammary fatpads (20). To determine if mGBP-2 expression correlated with better outcome in these cell lines, we first asked whether mGBP-2 was differential expressed in the two cell lines. While the non-metastatic 67NR cells expressed mGBP-2, the highly metastatic 4T1 cells expressed very low levels (Figure 2A). The documented migratory differences of these cells were confirmed (Figure 2B). mGBP-2 expression is inversely correlated with migration in these TNBC cells.

**Figure 2.**
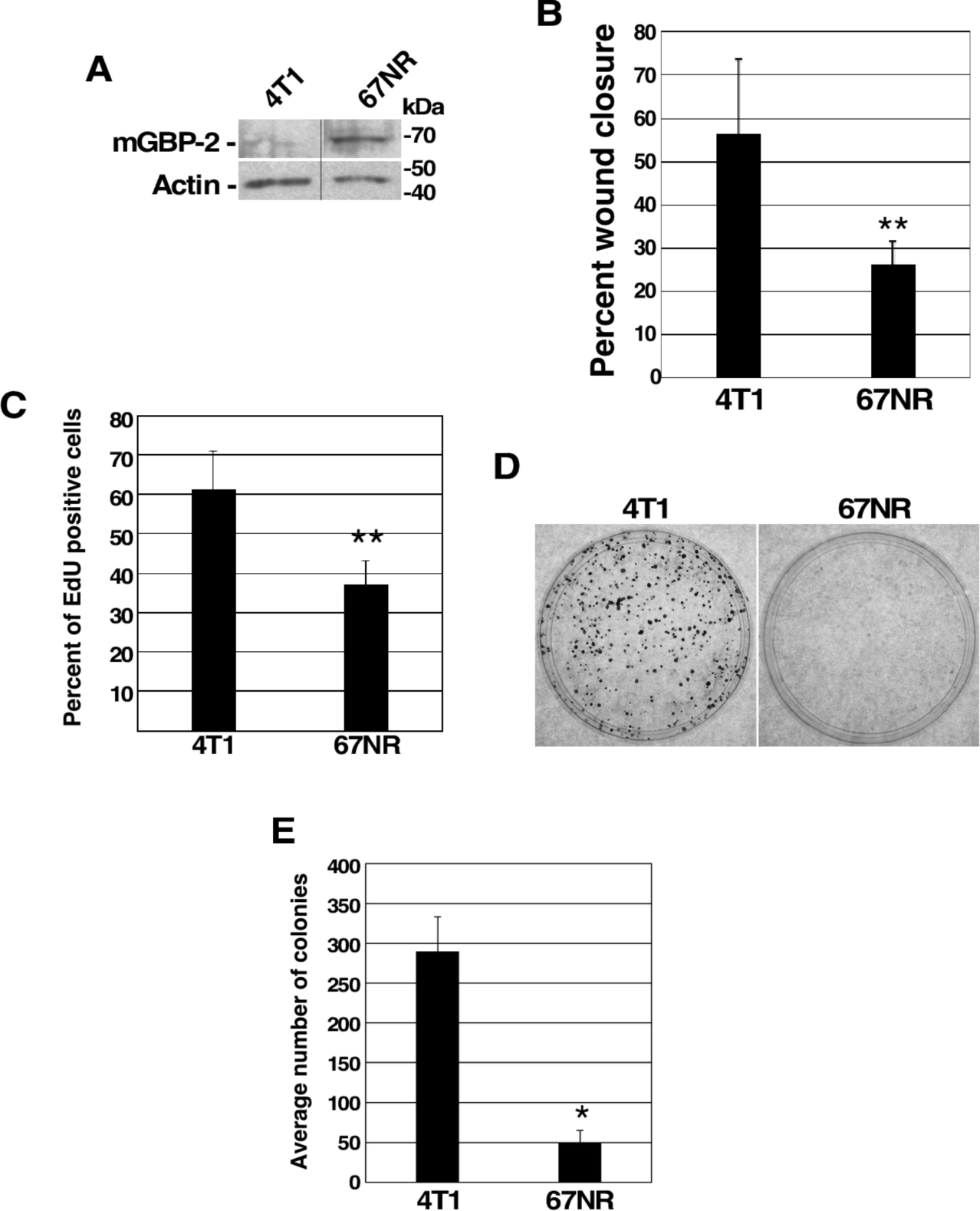
mGBP-2 expression inversely correlates with migration and proliferation in murine TNBC cell lines. **A).** Lysates from 4T1 and 67NR cells (20 μg) were analyzed for mGBP-2 and actin by western immunoblot (WB). A representative blot is shown (n = 3). **B).** Confluent 4T1 and 67NR cells were scratch with a 200 μl pipette tip and wound closure was analyzed as described in Methods. Wound closure was presented as average closure ± standard deviation (SD) (** = p < 0.01, n = 3). **C).** 4T1 (3 × 10^5^ cells/coverslip) and 67NR cells (4 × 10^5^ cells/coverslip) were cultured in duplicates in 6-well dishes and analyzed for S phase cells as described. The graph depicts the average percentage of EdU positive cells ± SD (** = p <0.01, n = 3). **D).** 4T1 and 67NR cells (1 × 10^3^ cells/dish) were seeded in triplicate in 6-cm dishes. After 6 days, cells were fixed and stained with crystal violet. Representative images are shown. **E).** All of the colonies with greater than 50 cells were counted per plate and represented as the average number of colonies per cell line ± SD (* = p <0.05, n = 3).

### mGBP-2 does not alter 4T1 or 67NR cell proliferation

The increased wound healing observed with 4T1 cells could reflect either increased migration, increased proliferation, or a combination of both. GBPs are well documented to alter cell proliferation both *in vitro* and *in vivo* (4,5,13,15,18,28-30). In fact, the expression of hGBP-1 in murine breast cancer cells inhibits their proliferation both *in vitro* and *in vivo* (5). The proliferation of 4T1 and 67NR cells were examined by EdU incorporation (Figure 2C). 4T1 cells incorporated EdU into about 60% of the cells within 1 hour, which was a little less than 2 times as many S-phase cells as for 67NR cells (Figure 2C). Moreover, these data are consistent with previous studies showing that about 60% of unsynchronized 4T1 cells are in S phase (31,32). Because some EdU incorporation could accompany DNA repair, the increased proliferation of 4T1 cells was confirmed by colony forming assays (Figure 2D). 4T1 cells grew significantly more colonies of 50 or greater cells than 67NR cells (Figure 2E). 67NR and 4T1 colonies exhibit significantly different cellular morphologies (Supplemental Figure 1A). 4T1 cells grow in compact, tightly associated colonies and 67NR cells tended to spread out. The ability of 67NR cells to spread out made the counting of colonies a little more difficult. To assure ourselves that our data was good, the crystal violet from the colonies was re-suspended in detergent and the optical densities of the dishes were measured (Supplemental Figure 1B). Together these studies show that 4T1 cells proliferate significantly faster than 67NR cells. They also suggest that the differences observed in the wound healing assays could indeed reflect some level of contribution from proliferation.

To determine if mGBP-2 inhibits breast cancer cell proliferation, 4T1 cells were treated with IFN-*γ* to induce the expression of mGBP-2 (Figure 3A) and the cells were examined for changes in proliferation (Figure 3B-D). Interferon treatment to increase mGBP-2 does not alter 4T1 proliferation as measured by EdU incorporation (Figure 3B) or colony formation (Figure 3D). Whether reducing the amount of mGBP-2 in 67NR cells would promote their proliferation was also examined (Figure 3E). 67NR cells were stably transfected with constructs containing either shRNAs against eGFP or shRNAs against mGBP-2 and stable pools of cells were isolated (Figure 3E). 67NR cells 3B and 3C showed greater than 90% knockdown of mGBP-2 (Figure 3E) and were used for subsequent experiments and designated KD#1 and KD #2 respectively. 67NR cells 2A with eGFP shRNA were designated as controls. These cells were examined for changes in proliferation (Figure 3F). Knocking down mGBP-2 in 67NR cells did not alter their proliferation. Together these data indicated that mGBP-2 does not inhibit the proliferation of these murine TNBC cells.

**Figure 3.**
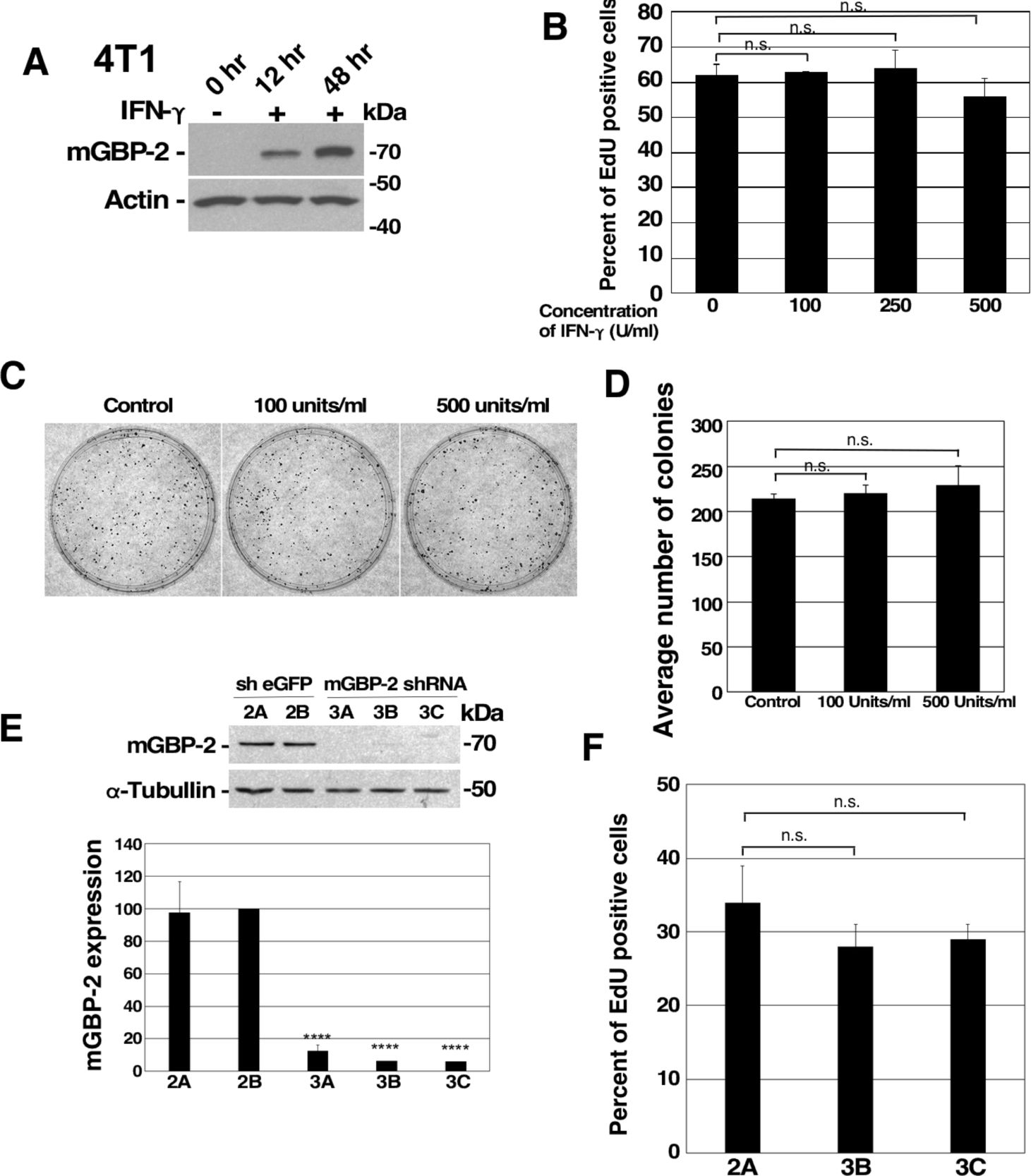
mGBP-2 does not inhibit 4T1 or 67NR cell proliferation. **A).** 4T1 cells (1 × 10^6^) were treated with 100 U/ml IFN-*γ* for the times indicated. Cells lysates (20 μg) were analyzed for mGBP-2 and actin by WB. A representative blot is shown (n = 3). **B).** 4T1 cells (1.5 × 10^4^ cells/coverslip) were treated with IFN-*γ* (0, 100, 250, and 500 U/ml). After 72 hrs, Click-it chemistry was performed as described in Materials and Methods. The graph depicts the average percentage of EdU positive cells ± SD (p = 0.7165, n = 3). **C).** 4T1 cells (1 × 10^3^ cells/dish) were treated with IFN-*γ* for 96 hrs. Cells were stained with crystal violet. Representative photomicrographs are shown**. D).** All of the colonies with 50 or more cells were counted per plate and represented as average number of colonies per condition ± SD (p = 0.3559, n = 2). **E).** Lysates (20 μg) from 67NR cells expressing mGBP-2 shRNA (3A, 3B, and 3C) or control shRNA (sh eGFP 2A and 2B) were analyzed for mGBP-2 and *α*-tubulin by WB. A representative gel is shown (n = 3). Data was analyzed as described and the ratio of mGBP-2 and *α*-tubulin densitometric values were represented as the average mGBP-2 expression ± S.D relative to control shRNA (shEGFP 2B) (**** = p < 0.0001, n = 3). **F).** 67NR cells containing sh eGFP 2A and two clones of mGBP-2 shRNA 3 (mGBP-2 shRNA 3B and mGBP-2 shRNA 3C) (3 × 10^5^ cells/coverslip) were cultured in duplicates in 6-well dishes analyzed for EdU incorporation as described. The graph depicts the average percentage of EdU positive cells ± SD (p = 0.0741, n = 3).

### mGBP-2 inhibits murine TNBC cell migration

GBPs, including mGBP-2, have been shown to alter cell migration and invasion (14,17,19,33-36). In particular, mGBP-2 inhibits cell spreading and migration of NIH 3T3 cells (19). Whether reducing the levels of mGBP-2 in the less migratory 67NR cells would lead to greater migration was examined using Boyden chambers. Greater than twice as many 67NR cells migrated when the level of mGBP-2 was reduced (Figure 4A). In addition, when the level of mGBP-2 was elevated in 4T1 cells by treatment with IFN-*γ* their migration was also significantly inhibited (Figure 4B). To determine whether mGBP-2 was the only IFN-*γ* induced protein responsible for the inhibition of migration, 4T1 cells stably expressing eGFP and mGBP-2 shRNAs were generated and treated with IFN-*γ* to induce mGBP-2 (Figure 4C). The migration of these cells was not influenced by the knockdown of mGBP-2 (Figure 4D), suggesting that mGBP-2 is not the only protein involved in the inhibition of cell migration by IFN-*γ*.

**Figure 4.**
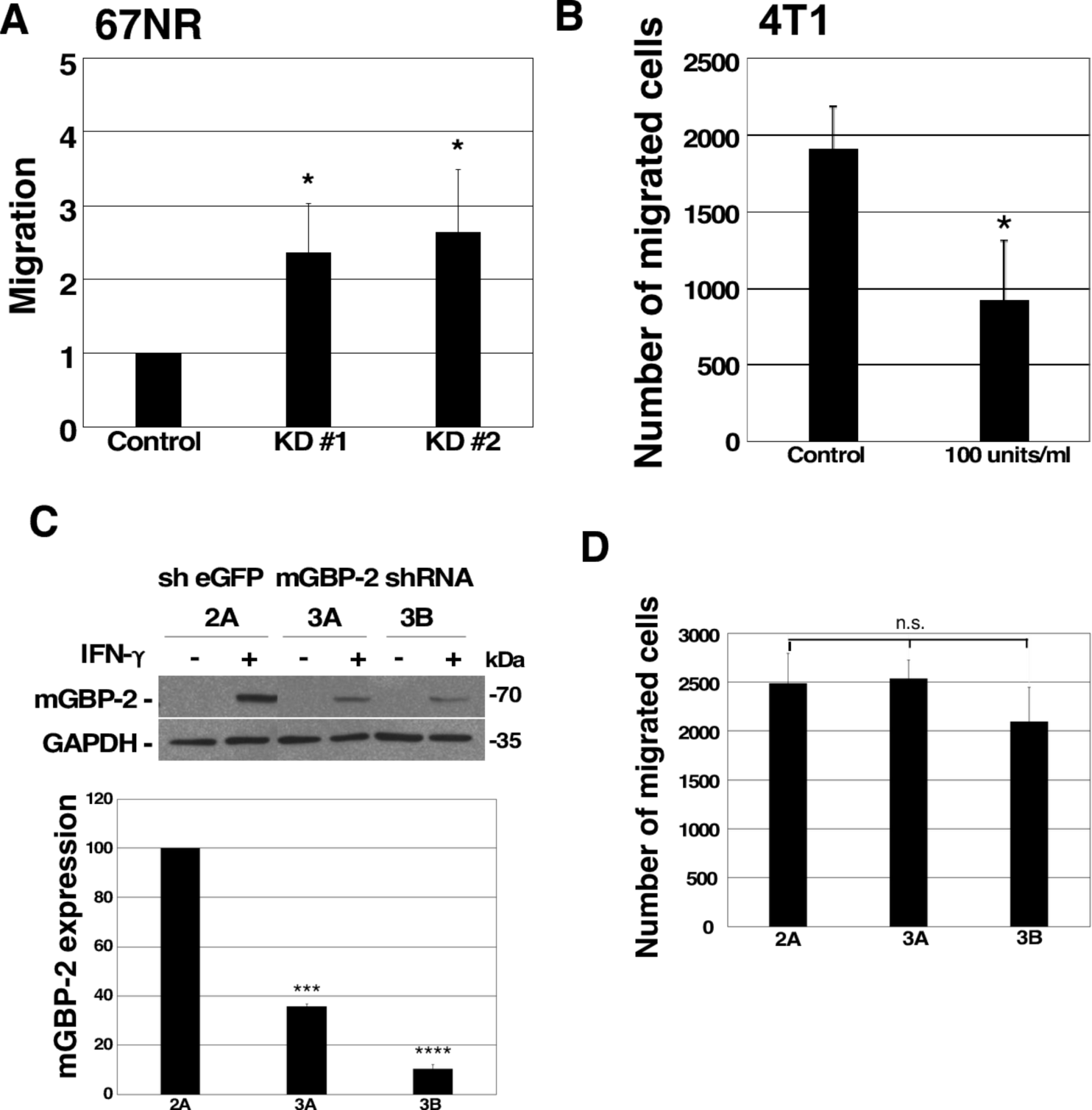
mGBP-2 inhibits 67NR cell migration but is not necessary for IFN-γ mediated inhibition of migration. **A).** Control, KD #1 and KD #2 67NR cells were seeded on Boyden chamber inserts coated with fibronectin. FBS (20%) in DMEM was added to the bottom well. Cells were allowed to migrate for 5 hrs. Membranes were processed and analyzed as described. All migrated cells were counted manually using ImageJ cell counter software. The graph presents the average migrated cells ± S.D. relative to control shRNA (sh eGFP 2A, which was assigned an arbitrary value of 1 (*p < 0.05, n = 3). The number of control cells that migrated ranged between 573 and 2167. **B).** 4T1 cells were pretreated with or without 100 U/ml IFN-*γ* for 24 hrs. They were then plated onto Boyden chambers in the presence or absence of IFN-*γ* and analyzed as described. The average number of migrated cells ± SD are shown (*p < 0.05, n = 3). **C).** Lysates from 4T1 cells expressing mGBP-2 shRNA or control shRNA (sh eGFP) and treated with 100 U/ml IFN-*γ* for 24 hrs, were analyzed for mGBP-2 and GAPDH. A representative blot is shown (n = 2). The ratio of mGBP-2 and GAPDH densitometric values were calculated and represented on the graph as the average mGBP-2 expression ± S.D relative to control shRNA (sh eGFP 2A), which was assigned an arbitrary value of 100 (*** = p < 0.001, **** = p < 0.0001, n = 2). **D).** 4T1 cells containing sh eGFP 2A, sh eGFP 2B, and two clones of mGBP-2 shRNA 3 (mGBP-2 shRNA 3A and mGBP-2 shRNA 3B) (5 × 10^4^) were pretreated with or without 100 U/ml IFN-*γ* for 24 hrs. The cells were plated onto Boyden chambers, allowed to migrate for 5 hours, and analyzed as described. All migrated cells were counted using ImageJ software. The average number of migrated cells ± SD are shown (p= 0.5775, n = 2).

### mGBP-2 alters 67NR actin cytoskeleton and morphology

GBPs have been documented to alter the actin cytoskeleton (12,16,19,35,37,38). Control and mGBP-2 KD 67NR cells were serum-starved and then activated by the addition of serum and the actin cytoskeleton and cellular morphology were analyzed (Figure 5A). On visual examination, the cells expressing mGBP-2 were more spread out, flatter, and appeared to contain more cell projections. The cells with reduced mGBP-2 appeared rounder. To confirm this, both the average number of cell projections greater the 10 µm per cell and the average projection length were determined. The control 67NR cells with mGBP-2 had greater than twice as many cell projections per cell than the KD cells (Figure 5B). In addition, their average projection lengths were about twice as long (Figure 5C). To quantify the more gross changes in cell morphology, the cells were scored as percent of cells having an elongation index greater than or equal to 2. Again, about twice as many of the control 67NR cells were elongated compared to the cells with lower mGBP-2 levels (Figure 5D). The observed morphology and shape changes indicate that mGBP-2 promotes breast cancer cell elongation upon serum activation. It also suggests that mGBP-2 may alter the activities of members of the Rho family of GTPases. Specifically, the finding of increased projections suggests an activation of CDC42, something never observed for a GBP.

**Figure 5.**
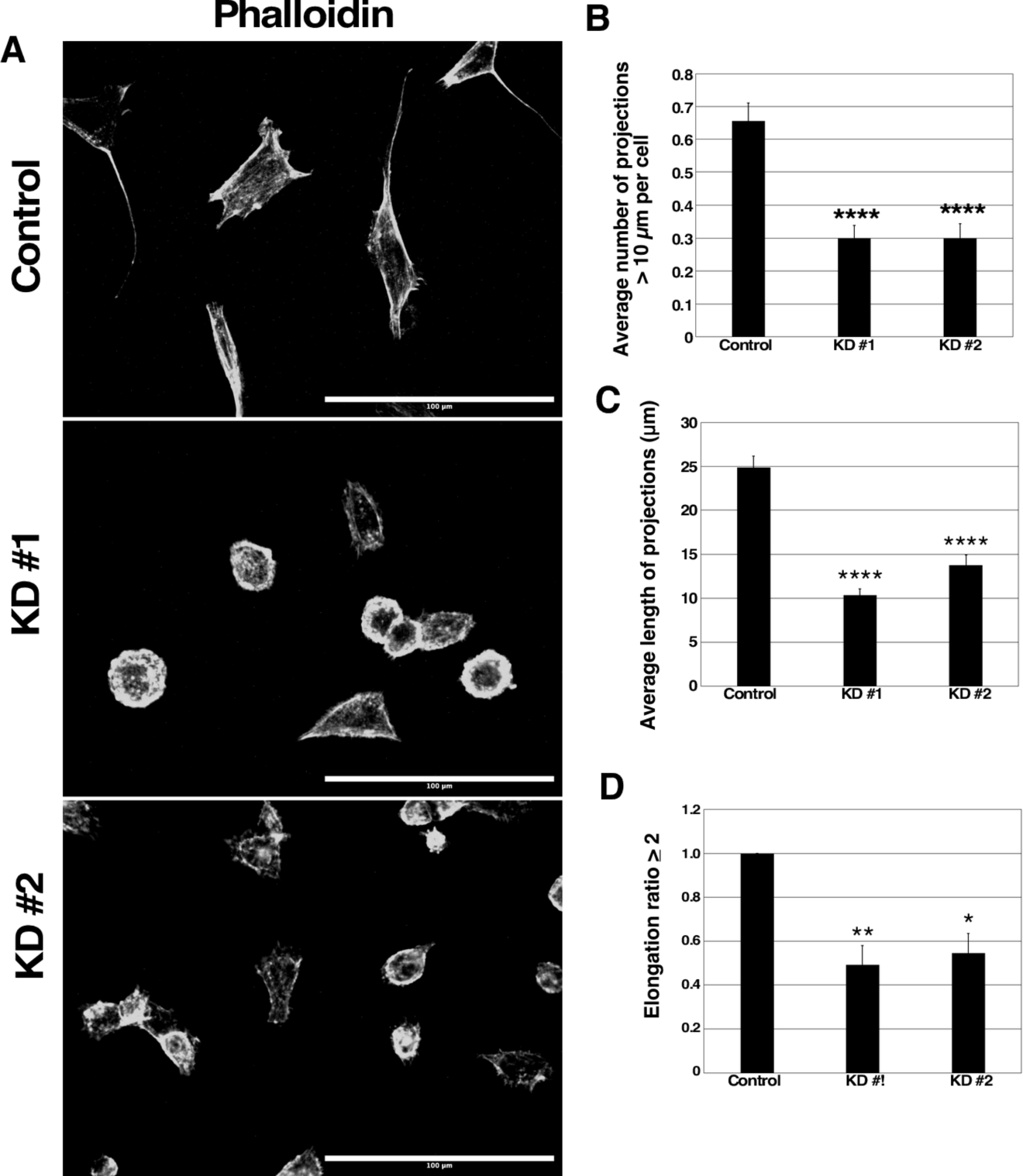
mGBP-2 alters the morphology of 67NR cells *in vitro*. **A).** Control, KD #1, and KD #2 67NR cells were serum starved for 3 hrs. Cells were then incubated with warm 20% FBS in DMEM for 30 mins. Cells were fixed, permeabilized, stained with Alexa fluor 594 phalloidin and 150 nM DAPI. Random fields were imaged on an EVOS FL Inverted Microscope at 40X using DAPI and Texas Red filters. Representative images are shown (Scale bars = 100 µm). **B).** The number of projections of 100 control, KD #1, and KD #2 67NR cells were counted. The graph represents the average number of projections per cell ± SEM (**** p < 0.0001, n=3). **C).** The length of projections of 100 control, KD #1, and KD #2 67NR cells were measured. The graph represents the average length of projections per cell ± SEM (**** p < 0.0001, n=3). **D).** The elongation ratios of 100 control, KD #1, and KD #2 67NR cells were measured. The graph depicts the average percentage of cells with an elongation >2 ± SEM (* p < 0.05, ** p < 0.01, n=3).

### mGBP-2 promotes the activation of CDC42 and RhoA and inhibits the activation of Rac1

To assess the effects of mGBP-2 on the activity of Rho GTPases, cells were serum starved and then activated with 20% FBS for 20 minutes. Rho activity levels were measured by effector pull-downs. The reduction of mGBP-2 in 67NR cells resulted in a 50% reduction in active CDC42 (Figure 6A). Therefore, mGBP-2 promotes the activation of CDC42 in 67NR cells. This is consistent with the finding of elevated numbers and lengths of projections in the 67NR cells expressing mGBP-2. The activity of GBPs on Rho A has also not been examined previously. mGBP-2 also modestly activates RhoA in 67NR cells (Figure 6B). mGBP-2 had previously been shown to inhibit the activation of Rac in NIH 3T3 cell fibroblasts downstream of plating on fibronectin, PDGF treatment, or TNF-*α* treatment (19,37). The reduction of mGBP-2 also resulted in a 15- to 20-fold increase in active Rac1, indicating that mGBP-2 also inhibits the activation of Rac1 in 67NR cells (Figure 6C).

**Figure 6.**
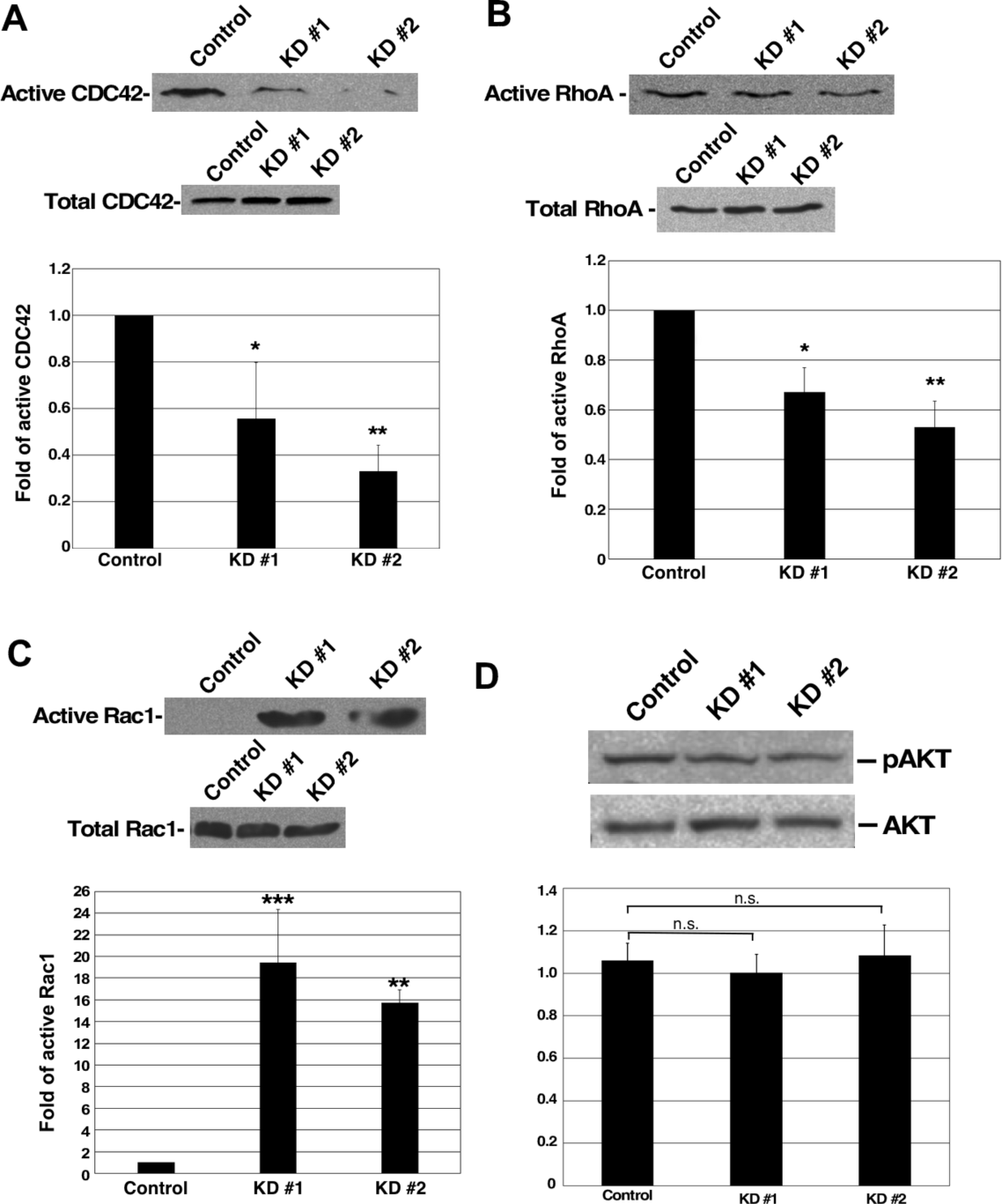
mGBP-2 promotes the activation of CDC42 and inhibits the activation of Rac1. **A).** Control, KD #1, and KD #2 67NR cells were serum starved for 12 hrs and then incubated with 20% FBS in DMEM for 30 mins, lysed, and analyzed for active CDC2 as described in Methods. The upper blot represents active CDC42 and the lower blot represents the total CDC42. Immunoblots from PBD pulldowns were quantified and results for levels of active CDC42 were normalized to total cellular CDC42 and then set to 1 for control 67NR cells (* = p < 0.05, ** = p < 0.01; n = 3**). B).** Control, KD #1, and KD #2 67NR cells were serum starved for 12 hrs and then incubated with 20% FBS in DMEM for 30 mins. Cell lysates were analyzed for active Rac1 as described in Methods. The upper blot represents active Rac1 and the lower blot represents the total Rac1. Immunoblots from PBD pulldowns were quantified and results for levels of active Rac1 were normalized to total cellular Rac1 and then set to 1 for control 67NR cells (** = p < 0.01, *** = p < 0.001; n = 3**). C).** Control, KD #1, and KD #2 67NR cells were serum starved for 12 hrs and then incubated with 20% FBS in DMEM for 30 mins. Cell lysates were analyzed for active RhoA as described in Methods. The upper blot represents active RhoA and the lower blot represents the total RhoA. Immunoblots from RBD pulldowns were quantified and results for levels of active RhoA were normalized to total cellular RhoA and then set to 1 for control 67NR (* = p < 0.05, ** = p < 0.01; n = 3). Representative blots are shown in A, B, and C. **D)** Control, KD #1, and KD #2 67NR cells were serum-starved for 18 hrs and then incubated with 20% FBS in DMEM for 30 min. Cell lysates were analyzed for phospho-Akt and total Akt. A representative blot is shown. Scanned X-ray films were uploaded into Image J to measure densitometric values of pAkt and total Akt. The ratio of pAkt and total Akt densitometric values were calculated and represented on the graph as the average pAkt ± SD relative to control, which was assigned an arbitrary value of 1 (p = 0.7617, n = 2).

### mGBP-2 inhibition of Rac is not accompanied by inhibition of activation of Akt

The inhibition of Rac by mGBP-2 in NIH 3T3 cells is accompanied by an almost complete inhibition of Akt activation downstream of plating on FN, as a consequence of inhibiting PI-3K (19). Interestingly, while mGBP-2 robustly inhibits Rac activation in 67NR cells, this inhibition is not accompanied by an inhibition of Akt activation (Figure 6D). This indicates that the inhibition of PI-3K may not be necessary for mGBP-2 to inhibit Rac in 67NR cells.

### mGBP-2 inhibits the generation of invadosomes in 67NR cells

In cultured cells, invasion is facilitated by specialized protrusions from the ventral surface of cells, known as invadopodia or podosomes. These structures degrade the ECM beneath cells to promote cell invasion, a process believed to be necessary for most cancer cell invasion (39,40). Podosomes and invadopodia have similar molecular components, morphologies, and functions and have been collectively referred to as invadosomes (41). While invadosomes can be formed by invasive cells in the absence of stimulation, this generally occurs at low frequency. A variety of growth factors induce invadosome formation, with their common features being activation of common signaling molecules such as Src, PI3-K, and the Rho family of GTPases (reviewed in (41,42)). For this study, we used phobol ester to promote invadopodia formation by activating Protein Kinase C (PKC) (42,43). Invadopodia can be recognized as small punctate or ring structures on the ventral surface of cells that co-stain for actin and cortactin (43,44). These are often found under or close to the nucleus. 4T1 cells have previously been shown to generate invadopodia (44). In the absence of growth factor or phorbol ester stimulation, 67NR cells did not express appreciable invadopodia, while 4T1 expressed invadopodia (as identified by co-staining of actin and cortactin) in just over 50% of the cells (44). In our hands, phorbol ester treatment of 4T1 cells also resulted in just over 50% of the cells making invadopodia (data not shown). However, phorbol ester treatment of 67NR cells resulted in low level of invadopodia formation, which was significantly increased when mGBP-2 expression was reduced (Figure 7).

**Figure 7.**
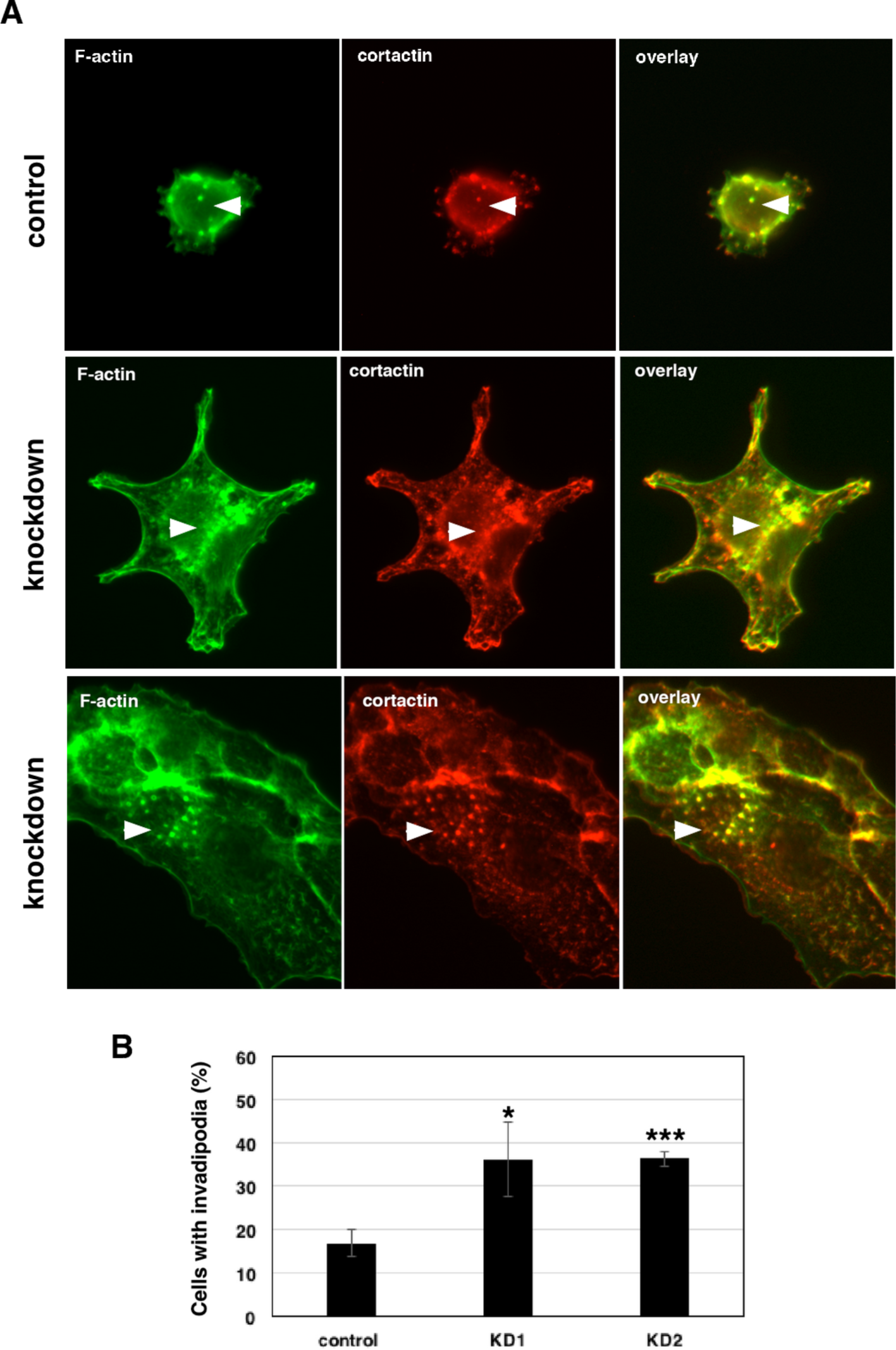
mGBP-2 inhibits invadopodia formation. Cells plated on coverslips and allowed to adhere overnight were treated with 1 µM PDBu for 30 minutes, fixed, and stained for cortactin, actin, and DAPI as described in Materials and Methods. **A).** Images of invadipodia in the cell lines. **B).** The percentage of cells containing invadipodia were determined for each cell type and represented as the mean ± SD (* = p < 0.05 and *** = p < 0.001; n = 3).

Together these data demonstrate that mGBP-2 acts to improve breast cancer prognosis by inhibiting migration and the assembly of intracellular structures needed for invasion.

## Discussion

Because GBP expression correlates with improved prognosis in breast cancers, this study specifically addressed whether GBP-2 contributes to cell autonomous changes that could result in improved prognosis. To answer this, the 4T1 model of murine triple-negative breast cancer was used (20). Both the 4T1 cells and 67NR cells came from the same spontaneously arising breast tumor. mGBP-2 was expressed in the non-metastatic, poorly migratory 67NR and not in the metastatic, highly migratory 4T1 cells (Figure 2). While GBPs, including mGBP-2, have been shown to modulate cell proliferation, mGBP-2 does not alter the proliferation of 4T1 or 67NR cells (Figure 3). What mGBP-2 does, that would be expected to improve breast cancer prognosis, is inhibit their migration and invadosome formation (Figure 4,7). Knocking down mGBP-2 expression in 67NR cells results in a significant increase in migration and invadopodia formation (Figure 4,7).

mGBP-2 inhibits cell migration by altering the activity of members of the Rho family of GTPases, master regulators of the actin cytoskeleton (45). The 67NR cells which express mGBP-2 were of a more mesenchymal appearance than the 4T1 cells, which grew in tightly associated colonies (data not shown). In addition, the 67NR cells had more projections/filopodia (Figure 4). The presence of cell projections/filopodia suggested that in the presence of mGBP-2 the Rho family member, CDC 42, was activated. Consistent with this morphology, there was 40-70% more active CDC 42 than in the absence of mGBP-2 (Figure 5A). RhoA was also activated in the presence of mGBP-2 (Figure 5B). Interestingly, when mGBP-2 levels were significantly reduced in 67NR cells, the morphology became rounder (Figure 4D) and the presence of lamellipodia were more common (Figure 4A). These are features associated with activation of Rac1 (45). Consistent with this, Rac1 activity was lost in the presence of mGBP-2 but when mGBP-2 was absent was robustly activated (Figure 5C).

This is the first observation that a GBP can influence cell invasion by directly inhibiting the formation of invadosomes. mGBP-2 alters the activities of members of the Rho family of GTPases (Figure 6;(19)) and to inhibit PI3-K activation downsteam of integrin engagement (19). In 67NR cells, mGBP-2 did not inhibit P13-K activation by serum but it did modulate the activity of Rac1, Cdc42, and RhoA (Figure 6). Future studies will determine the molecular mechanisms by which mGBP-2/GBP-2 regulate Rho GTPases to inhibit migration/invasion and contribute to improved prognosis in breast cancer.

## Authors’ Contributions

Conception and design: D.J. Vestal, G. Nyabuto

Development of methodology: G. Nyabuto, J. Wilson

Acquisition of data: G. Nyabuto, J. Wilson, A. Abnave, S. Heilman, R. Kalb

Writing, review, and/or revisions of the manuscript: D.J. Vestal, G. Nyabuto

Study supervision: D.J. Vestal

## Acknowledgements

The work was supported by the University of Toledo. The authors would like to acknowledge the gift of reagents and expertise on Rho family protein activity assays from Dr. Rafael Garcia-Mata, suggestions on Boyden chambers for Dr. Kathryn Eisenmann and Krista Peatee, and instruction on the Cytation 5 from Drs. Andrea Kalinoski and David Weaver.

---

**Supplemental Figure 1.**
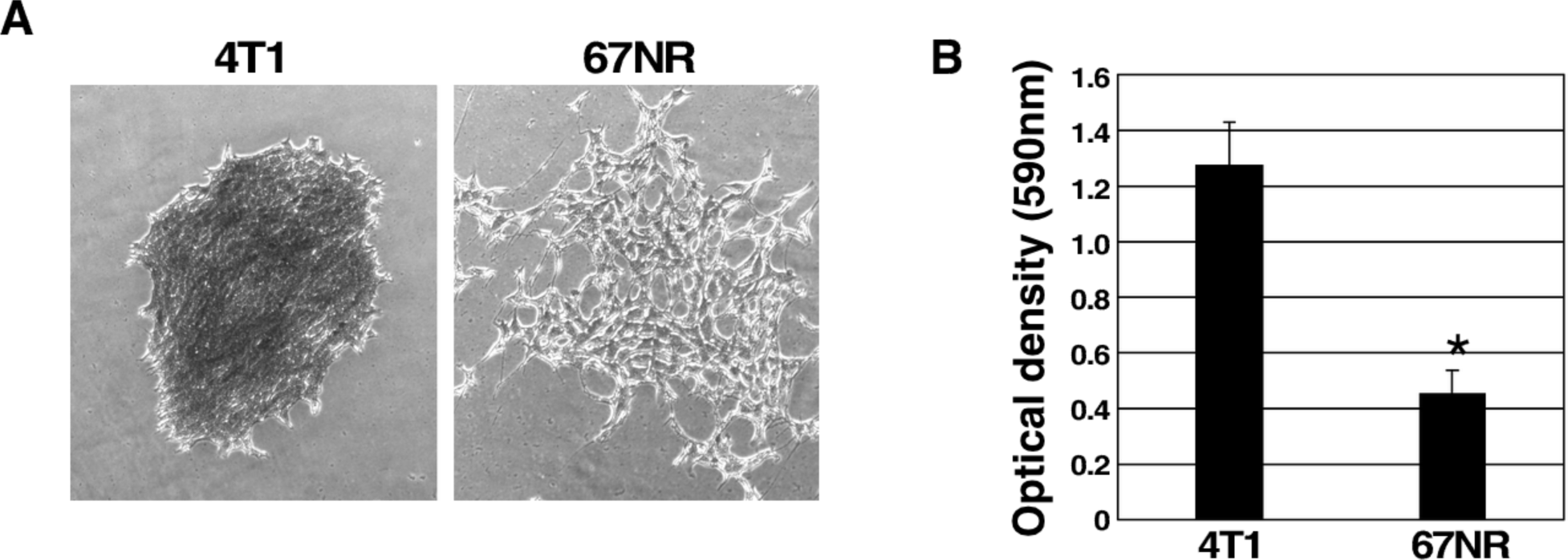
4T1 cells have more proliferating cells than 67NR cells. **A).** Representative photomicrographs (40X) of a single colony of 4T1 and 67NR cells show differences in colony morphology. **B).** Crystal violet was extracted from the plates in part D with 1% SDS and quantified at 590 nm. The average optical densities ± SD are shown (**\*** = p <0.05, n = 3).

## The uncropped autorads for the Western blots follow

**Fig. 2A.**
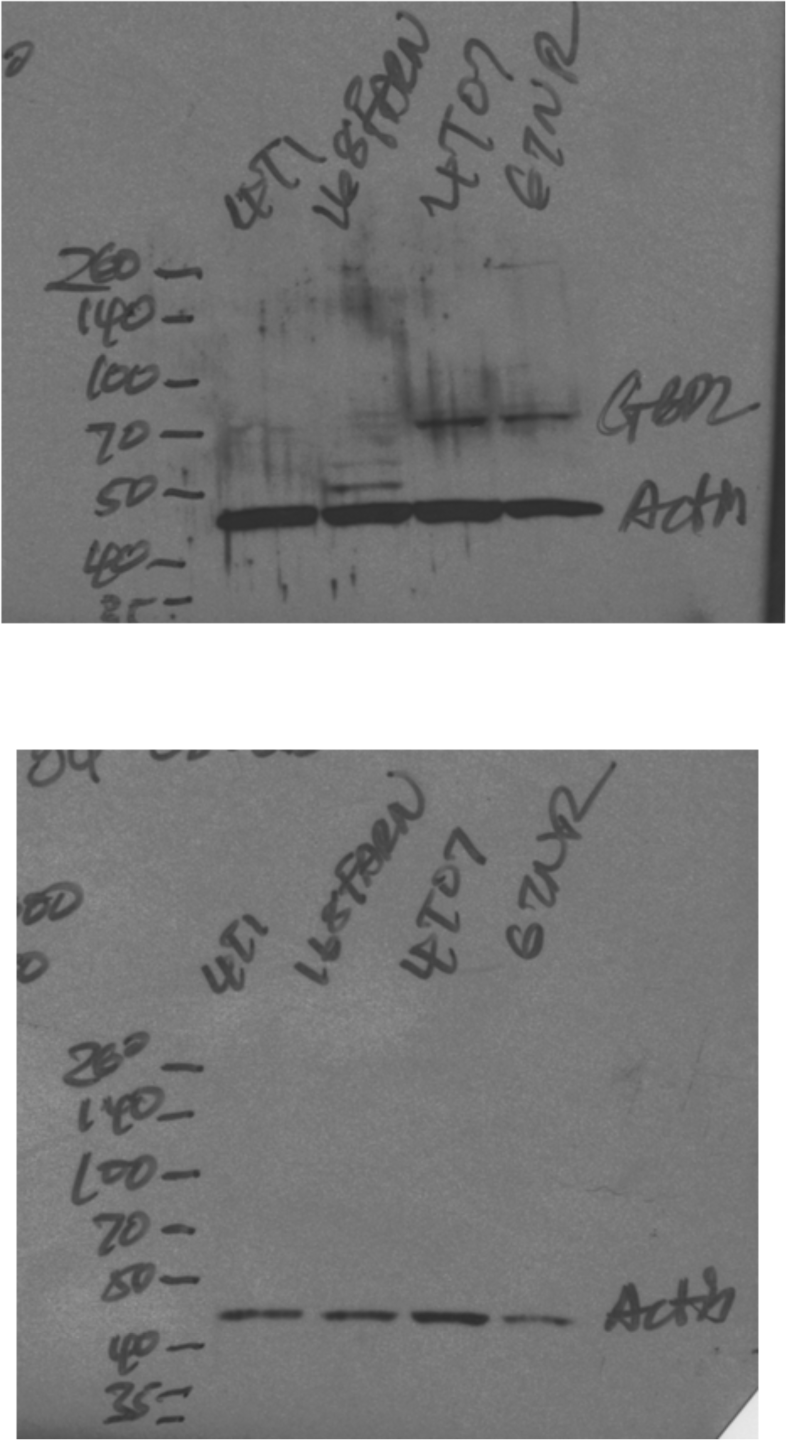

**Fig. 3A.**
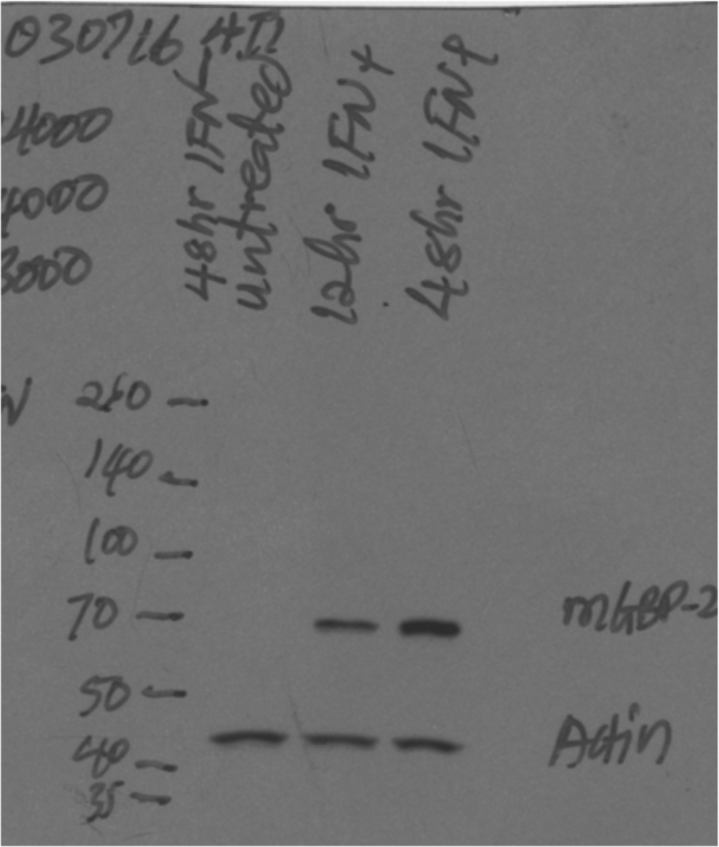

**Fig. 3E.**
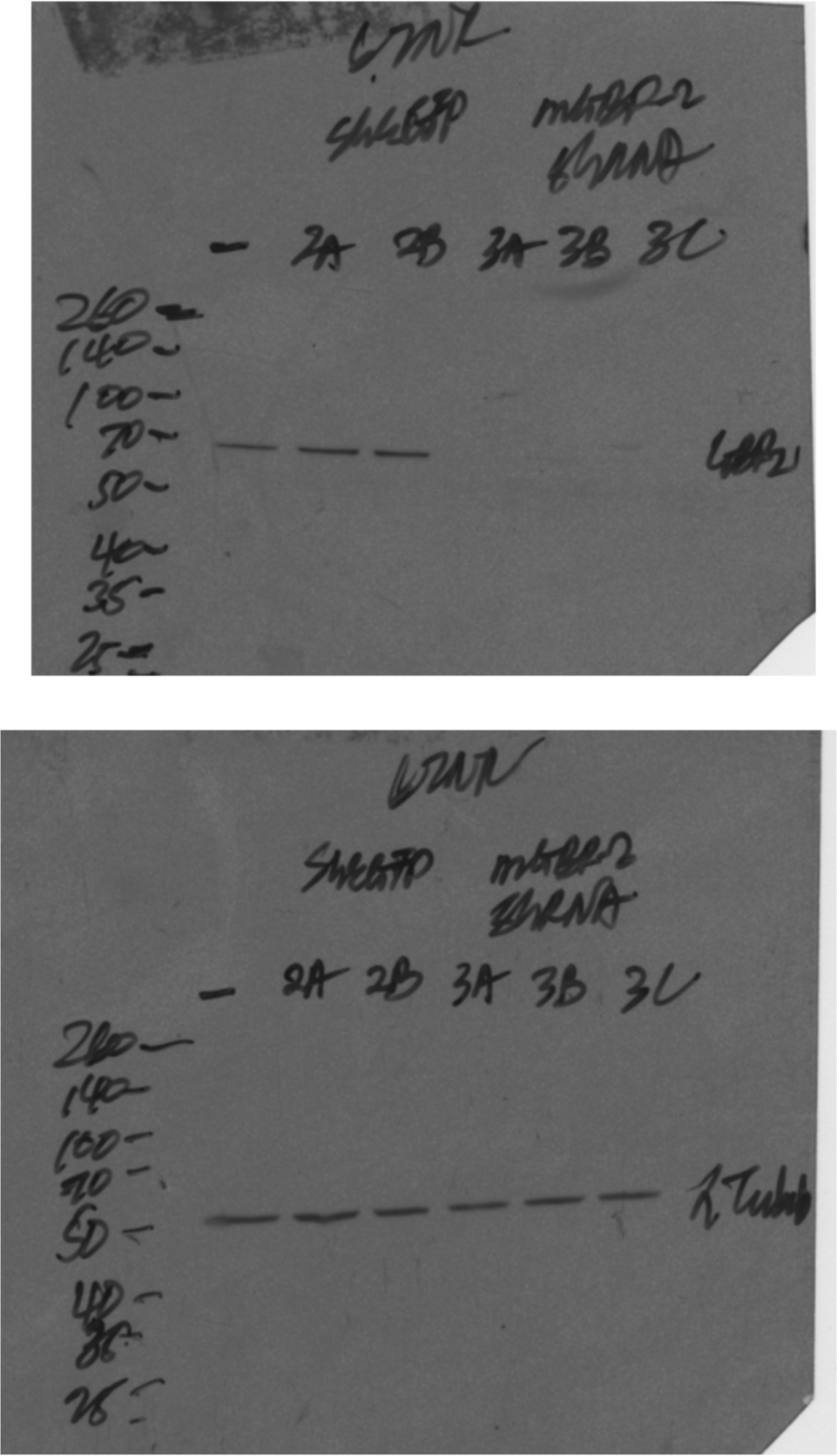

**Fig. 4C.**
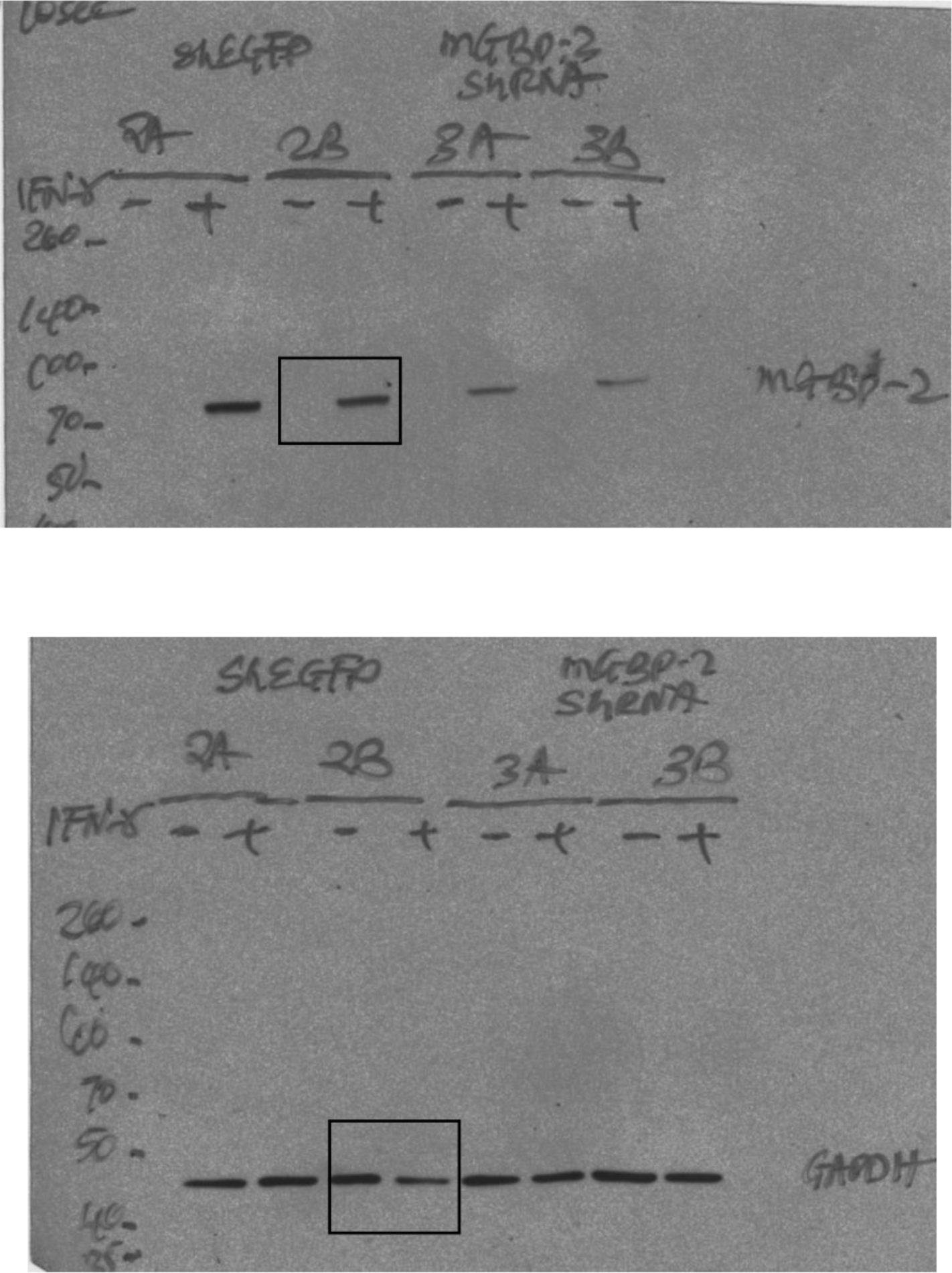

The area within the black box was cropped out of the figure.

**Fig. 6A.**
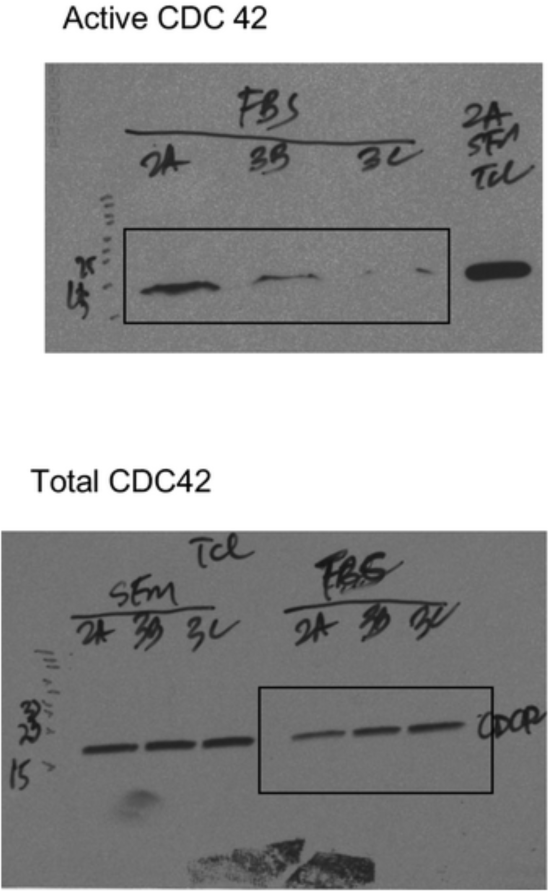

**Fig. 6B.**
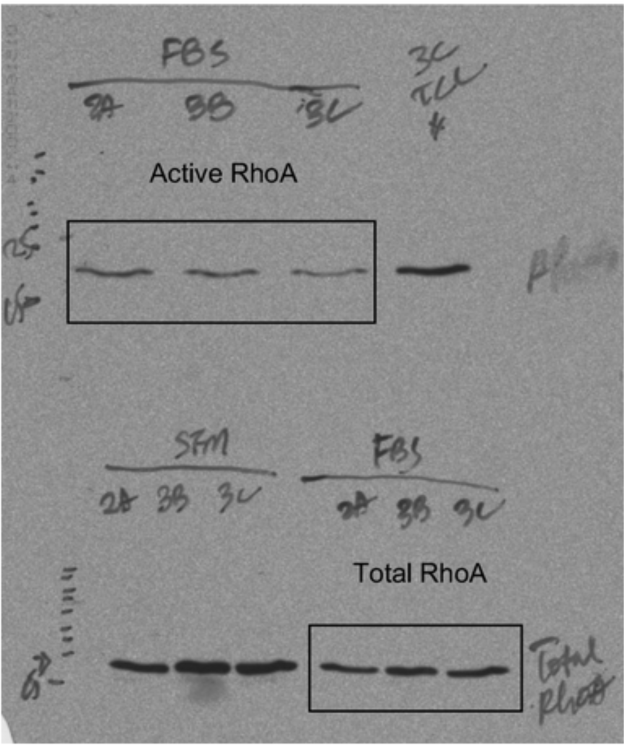

**Fig. 6C.**
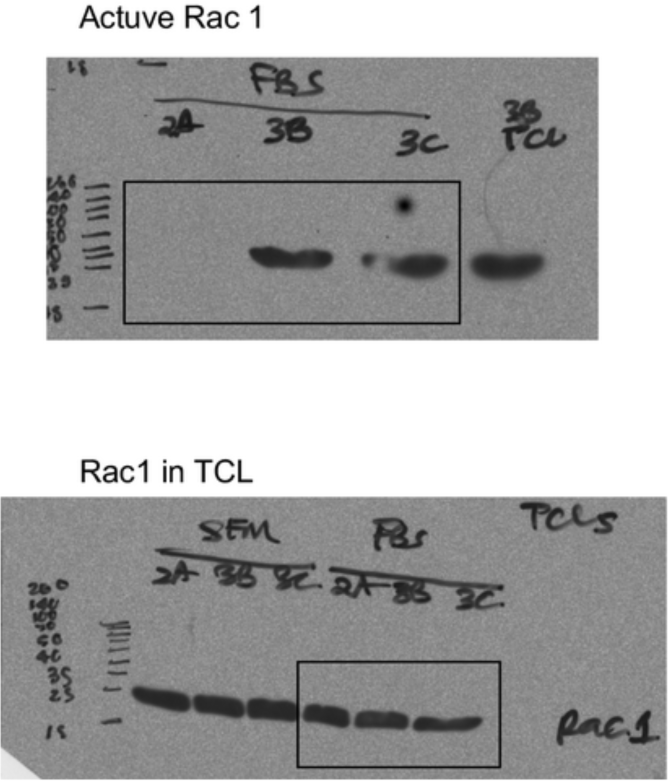

**Fig. 6D.**
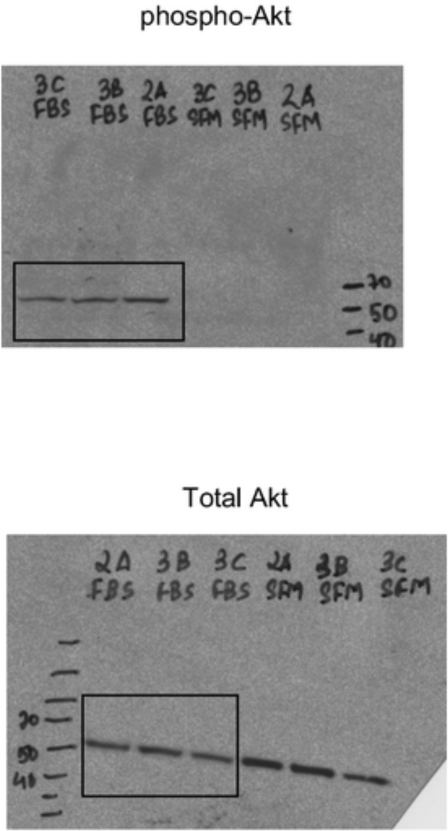

The regions within the black boxes are those in the figures.

